# ProHap enables proteomic database generation accounting for population diversity

**DOI:** 10.1101/2023.12.24.572591

**Authors:** Jakub Vašíček, Ksenia G. Kuznetsova, Dafni Skiadopoulou, Pål R. Njølstad, Stefan Johansson, Stefan Bruckner, Lukas Käll, Marc Vaudel

## Abstract

Amid the advances in genomics, the availability of large reference panels of human haplotypes is key to account for human diversity within and across populations. However, mass spectrometry-based proteomics does not benefit from this information. To address this gap, we introduce ProHap, a Python-based tool that constructs protein sequence databases from phased genotypes of reference panels. ProHap empowers researchers to account for haplotypic diversity in proteomic searches.

## Main

As population-wide genetic data sets and fine charting of the genome become available [1], our ability to map haplotypes within and across populations has expanded dramatically. This has allowed for the imputation of previously unobserved genotypes, increasing the power of genome-wide association studies [2]. However, mass spectrometry-based analysis of proteins has, thus far, not accounted for these crucial facets of human diversity. This oversight obscures a portion of the proteome, introducing an unintended bias against populations not well represented by reference sequences. We propose to address this challenge by proteogenomics, a field integrating proteomics and genomics in a joint approach to capture the consequences of genetic variation in proteins [3], [4].

In proteogenomics, custom databases of protein sequences are generated to allow for the identification of alternative peptide sequences in data sets generated by tandem mass spectrometry (MS/MS). Tools generating these protein databases are typically based on the *in silico* translation of custom RNA sequences (such as *customProDB* [5]), or on aligning genetic variants with spliced transcript sequences provided by Ensembl before the translation (such as *pgdb* from *py-pgatk* [6]). However, these tools do not allow the generation of entire protein haplotypes from human reference panels, and therefore the databases do not reflect the sequences found in human populations. With Haplosaurus, Spooner et al. [7] enabled inspecting the influence of haplotypes on proteins. As Haplosaurus is tailored for the examination of specific genes primarily focusing on drug design [7], it is not suited for the generation of proteome-wide sequence databases.

Here, we present ProHap — a lightweight bioinformatic tool designed to efficiently construct protein sequence databases from large reference panels of human haplotypes (Figure 1). ProHap also allows the inclusion of disease-specific rare variants, giving researchers the opportunity to create protein sequence databases capturing the effect of both common and rare variation. The database generated by ProHap can be matched by proteomic search engines, improving the coverage of the proteome varying within and across populations. Subsequently, ProHap enables the mapping of identified peptides to the genes and transcripts possibly encoding them, hence allowing the evaluation of the coverage of splicing sites, protein haplotypes, and alleles of interest. ProHap is available as a Snakemake pipeline under a permissive open source license (MIT [8]), and features a web-based graphical interface to generate the configuration file.

**Figure 1:**
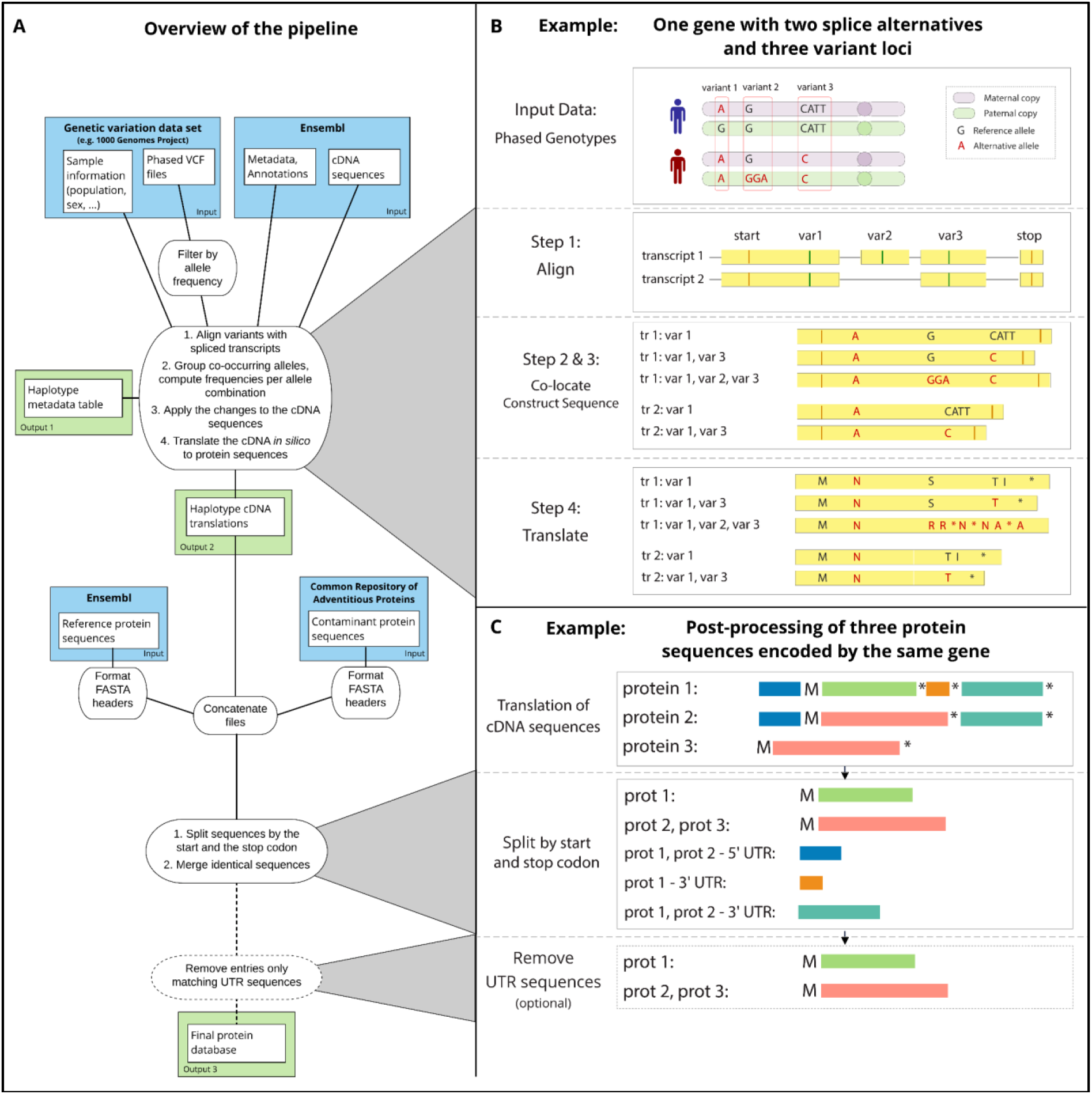
A: Diagram detailing the workflow of ProHap. Input files are highlighted in blue, output files highlighted in green. B: Construction and *in silico* translation of haplotype cDNA sequences. C: Optimizing the protein sequence database: splitting the sequences by translation events (start and stop codons), merging the overlapping sequences, optionally removing sequences matching translations of the UTR regions. The M letter stands for the initial methionine in the protein sequence, the asterisk (*) represents a stop codon.

To demonstrate the use of ProHap, we have constructed a reference database of protein haplotype sequences from the phased genotypes of the 1000 Genomes Project [9], build GRCh38 of the reference genome [10]. The 1000 Genomes Project provides a clustering of participants into populations, themselves pooled into five ancestrally related groups [9], referred to as superpopulations: *African, American, East Asian, European*, and *South Asian*. We have built protein haplotypes for every superpopulation. The resulting databases, available as supplementary information, account for common alleles with a frequency of 1% or higher and exclude rare variation. Moreover, we have built a complete database using the genotypes of all the participants of the 1000 Genomes project, which contains the alternative allele for 56,242 non-synonymous variants, distributed across 254,143 unique protein sequences. Among these variants, 52,361 represent single amino acid polymorphisms (SAPs), while 128 introduce stop codon alterations (stop-lost variants). Additionally, 884 variants manifest as in-frame insertions or deletions (indels), and 461 are frameshift variants. Furthermore, the haplotypes include the alternative allele for 1,606 variants always located at splicing sites, where the prediction of the protein sequence is challenging, and for 32,659 variants that are synonymous in all considered splicing alternatives, and therefore cannot be considered as discoverable in protein sequences.

Within the database, certain variants exhibit context-dependent consequences depending upon the splicing of the associated genes. Notably, 775 variants may give rise to either synonymous mutations or SAPs depending on the transcript considered. Additionally, 12 variants have the potential to cause either a loss of the stop codon or introduce a SAP, while 10 variants cause either a loss of the stop codon or a synonymous mutation. Furthermore, we identified 213 non-synonymous and non-frameshift variants that may occur in haplotypes following a frameshift, adding a layer of complexity to their interpretation. Of these variants, 119 occur exclusively between a frameshift and the subsequent early stop codon in all haplotypes, rendering them indiscernible unless the entire haplotype is considered.

To assess the discoverability of these variants in peptides, we conducted an *in silico* digestion using six different proteases: trypsin, Lys-C, Lys-N, Glu-C, Asp-N allowing for up to two missed cleavages, and chymotrypsin allowing for up to four missed cleavages. We found that for 54,679 variants, the alternative allele can be detected in peptides using at least one of the six proteases. The discoverability rates varied by consequence type, with 98.2% of SAPs, 93.6% of in-frame indels, 94.9% of stop-lost variants, and 99.1% of frameshift variants being detectable.

We classify the peptides containing the consequences of variation into three categories: (1) Single-variant peptides carry the consequence of one genetic variant, remaining unaffected by other variants in linkage disequilibrium (LD); (2) Multi-variant peptides encompass the consequences of at least two variants in LD [11]; and (3) Frameshift peptides are mapped to regions inclusive or downstream of a frameshift variant, regardless of whether they contain additional variant consequences.

The analysis revealed that 82% of variants are discoverable solely in single-variant peptides, while 14.9% can be detected in both single-variant and multi-variant peptides, depending upon the number of missed cleavage sites and the proteases utilized. A smaller proportion, 1.3%, is exclusively discoverable in multi-variant peptides. Moreover, 1.5% of variants are potentially discoverable in frameshift peptides, and 1% are detectable in frameshift peptides exclusively. These findings underscore the ability of ProHap to capture the consequence of a broad range of genetic variation that was so far not available to proteomics researchers. Overall, we observe that variant peptides account for up to 14.1% of all encoded sequences. However, the variation is unevenly distributed among the superpopulations included in the 1000 Genomes Project panel (Figure 2A).

**Figure 2:**
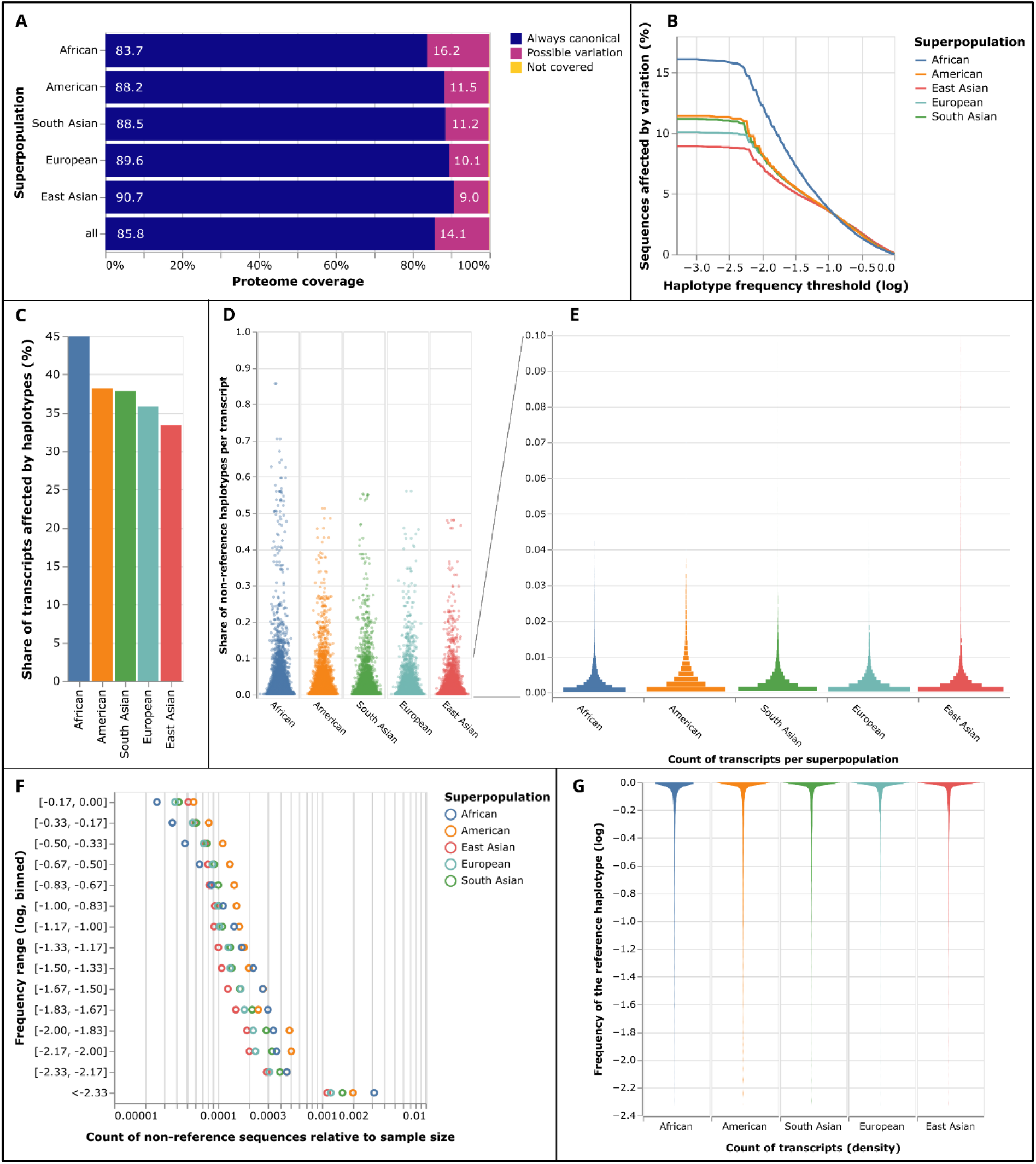
A: Proteome coverage expressed in terms of the percentage of amino acids considering all possible splicing alternatives of all genes. I.e., if 7% of the proteome possibly contains variation, 7 out of 100 residues belong to at least one discoverable peptide containing the product of a variant. B: Percentage of the proteome affected by variation dependent on the lower haplotype frequency threshold. C: Percentage of transcripts affected by haplotypic variation per superpopulation. D: Count of haplotype sequences observed per transcript, relative to the maximal possible number of sequences (double the count of samples in each group). Each point represents a transcript. E: Count of transcripts (x-axis) observed to produce a given number of haplotype sequences (y-axis). Transcripts unaffected by haplotypes in any population are not considered. F: Count of unique protein sequences observed in each frequency interval per superpopulation, relative to the maximal count of unique sequences (double the count of samples in each group * total count of transcripts). G: Violin plot showing the distribution of the frequencies of the reference haplotype for each transcript per superpopulation. Transcripts unaffected by haplotypes in any population are not considered.

As per one of the primary objectives of the 1000 Genomes Project — to encompass diverse populations globally — it was observed that participants assigned to the African superpopulation exhibited the highest numbers of variants likely to impact gene function [9]. Correspondingly, our analysis using ProHap reveals a notable increase in the proportion of the proteome affected by common haplotypes (Figure 2A, 2B, 2C), an increase in the count of unique haplotype sequences per spliced transcript (Figure 2D, 2E) for participants assigned to the African superpopulation, and an increase in the coverall count of non-canonical protein sequences (Figure 2F) for participants assigned to the African and American superpopulations. Concurrently, there is a decrease in the frequency of the reference haplotype per spliced transcript in the African superpopulation (Figure 2G). When analysing diverse samples, the databases generated by ProHap will therefore represent the proteome of the participants more comprehensively and fairly than the reference proteome, enhancing our insight into the impact of genetic differences on protein sequences.

ProHap is a Python-based tool built to efficiently generate protein haplotype databases from phased genotypes of reference population panels. This makes it possible to comprehensively consider different types of genetic variants: single nucleotide polymorphisms (SNPs), insertions, deletions, alterations of the stop codon, and frameshifts, and to account for population diversity in proteomic studies. The link between identified variants in peptides and their genetic origins bridges the gap between genomic and proteomic studies, which will enable a deeper understanding of genetic influences on protein sequences. ProHap’s modular design allows users to generate updated databases aligned with the latest genome build and Ensembl annotation. Implemented as a Snakemake pipeline, it ensures transparent and reproducible outcomes. The tool offers a number of parameters to control database size, e.g., excluding sequences like untranslated regions (UTRs) of transcripts.

Crucially, as global genomic panels only partially capture the genetic diversity of underserved populations [12], ProHap’s adaptability to any phased genotype panel enables proteomic studies tailored to any specific population. Moreover, as a standalone pipeline, it can be executed on secure servers without relying on online genomic variation resources, thereby maintaining the ownership of and access to genomic data, which are crucial for equitable and inclusive research [13], [14]. Finally, as the study of the proteome in diverse populations emerges, we would like to stress the importance of carefully considering associated ethical implications to prevent misinterpretation, biases, and discrimination [15].

## Methods

### Public data sources

We compute the protein haplotypes in *ProHap* based on a VCF (variant call format) file of phased genotypes (Figure 1A). The data set is expected to come as a separate VCF file for each chromosome, and the rows in each file are expected to be sorted by the respective variant’s genomic location, 5’-most to 3’-most. Such data is made available by the 1000 Genomes Project [9], and the updated version contains phased genotypes of 2,709 samples from populations worldwide aligned with the GRCh38 reference genome [10]. Participants of the 1000 Genomes Project were assigned into populations, which were further pooled into five ancestrally related groups [9], referred to as superpopulations. Table 1 shows the number of samples in each superpopulation which are available in the VCF files.

**Table 1:** Number of participants per superpopulation in the 1000 Genomes Project data set.

| Superpopulation | Number of samples |
| --- | --- |
| European | 522 |
| East Asian | 515 |
| American | 348 |
| South Asian | 492 |
| African | 671 |
| All | 2,548 |

The simplified version of the tool, *ProVar*, accepts one VCF file containing all the considered variants, ordered by chromosome and location within the chromosome, 5’-most to 3’-most. Please refer to the documentation (see below) for further details. While it is possible to use public data sets of common variation with *ProVar*, it is mainly meant for appending rare variation to the database of common haplotypes.

The variants in the VCF file are aligned with spliced transcripts provided by Ensembl [16]. Therefore, we use the Ensembl annotation file in the GTF format, as well as the databases of canonical cDNA and protein sequences. For the results demonstrated here, we used Ensembl version 110 [16].

### Database of protein sequences

The database of protein sequences is constructed in four steps, as illustrated in Figure 3. In the first step, the chromosome is scanned using a sweep line approach to assign variants (Figure 3A) to each possible splicing of the overlapping gene(s) (Figure 3B). Variants that do not intersect an exon or do not pass the allele frequency threshold are discarded. Next, for each included splicing of each gene, we store the combinations of variants for which the alternative alleles were observed co-occurring in the data set, along with the frequencies of occurrence of these combinations. Using these combinations of alleles, we construct the respective cDNA sequences (Figure 3C) and obtain the protein sequences by *in silico* translation (Figure 3D). Where available, the location of the main open reading frame (mORF) is identified from the annotation of the start and stop codons in Ensembl. If the annotation of the start codon is lacking, the mORF is assumed from the start of the transcript until the first annotated stop codon is reached. We provide options to perform the three-frame *in silico* translation in this case, or to skip the transcript completely.

**Figure 3:**
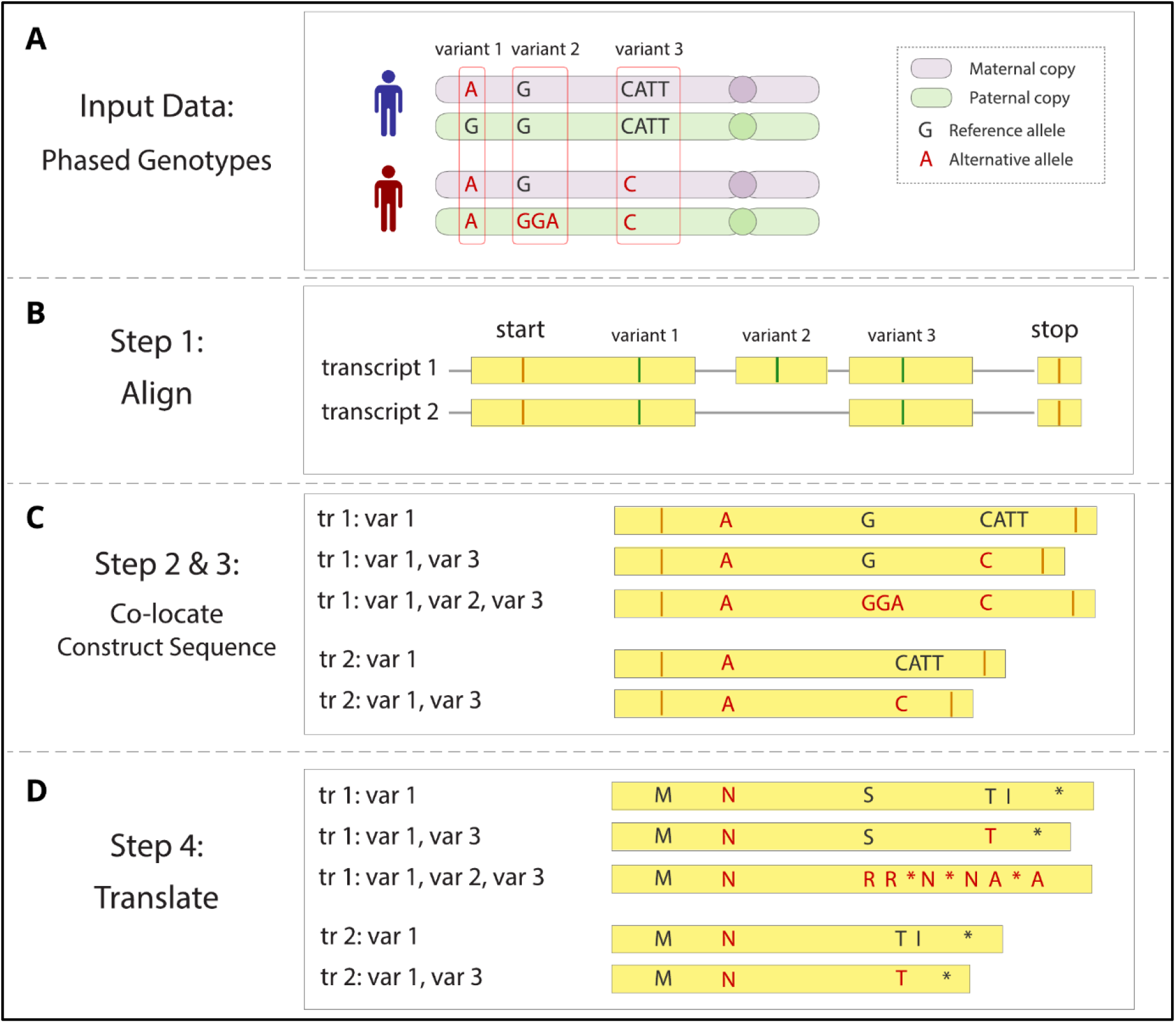
Construction of the database of protein haplotypes in four steps. A: input data consist of the variant call format (VCF) file of phased genotypes; B: the variants are aligned with two transcript sequences provided by Ensembl; C: variants where the alternative alleles co-occur are grouped per transcript and the alternative cDNA sequences are constructed for each such haplotype; D: cDNA sequences are translated *in silico* to obtain protein sequences, the asterisk (*) represents a stop codon.

Consequently, the translated sequences are split by the annotated start and stop codon locations, or by the observed stop codon locations in case of three-frame translation (Figure 4A). Identical sequences after splitting are then merged into the same FASTA entry (Figure 4B). Optionally, sequences matching only translations of untranslated regions (UTRs) are removed (Figure 4C).

**Figure 4:**
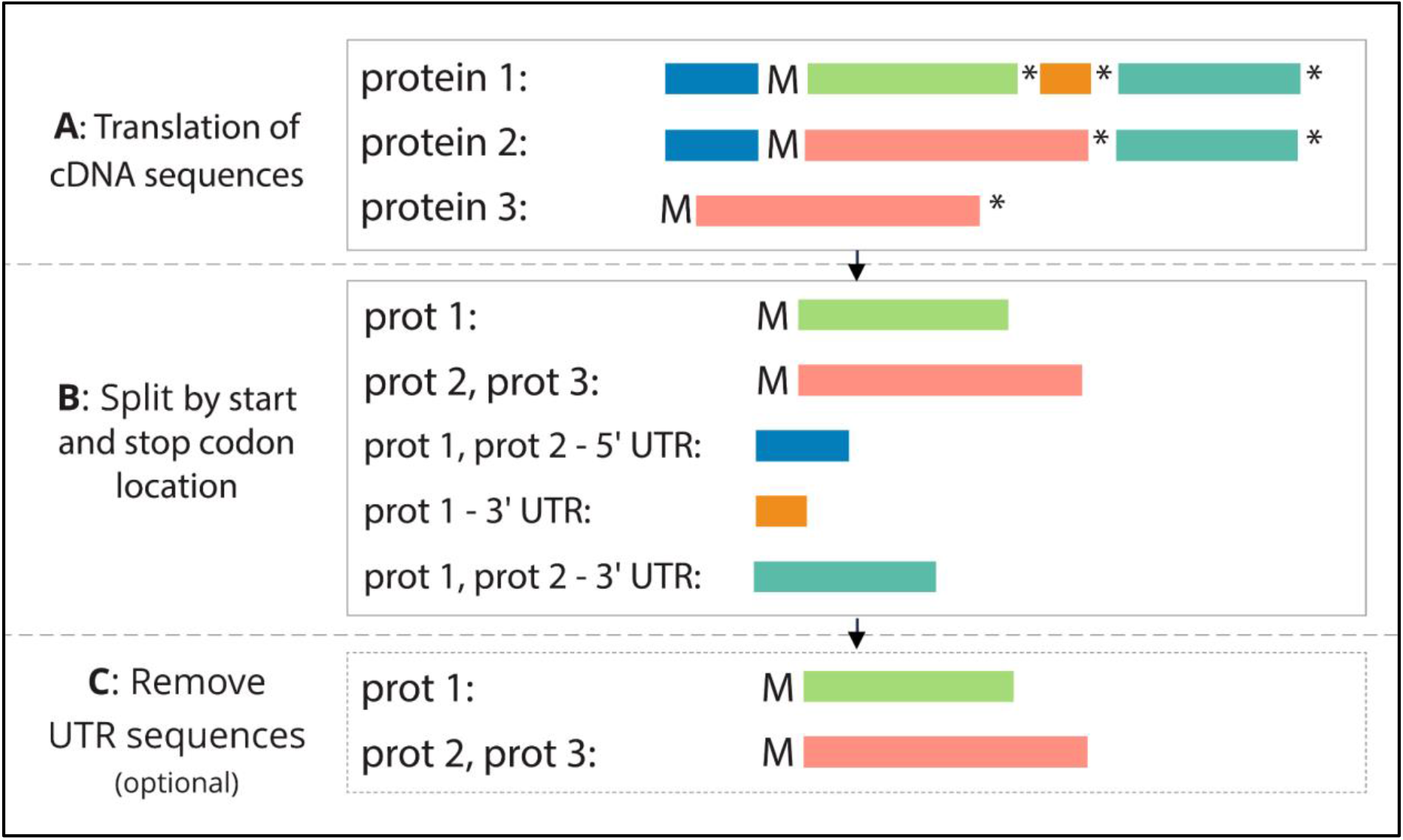
Post-processing steps on the database of protein sequences. The full translations of cDNA sequences (A) are split by the start and stop codon locations, and identical entries are merged (B). Optionally, sequences matching to translations of UTRs are removed (C). The M letter stands for the initial methionine in the protein sequence, the asterisk (*) represents a stop codon.

The result of the ProHap pipeline consists of three output files:

1. Haplotype table: Associated information for all the haplotypes included. This includes the combination of alleles, the protein consequences, frequency of occurrence, and other metadata (see documentation for details),
2. Haplotype FASTA: Database of all haplotype sequences prior to any merging or optimization in the FASTA format,
3. Optimized FASTA: Final protein sequence database in the FASTA format including the canonical proteome, contaminant sequences, and haplotype and/or variant sequences, optimized for use with SearchGUI [17].

### Quality control

We apply a set of quality control checks before the translation of the cDNA sequences to ensure the validity of the results. Namely, there are haplotypes which present conflicting alleles occurring at the same genetic location. We consider these cases to be errors in phasing and remove the samples where such conflicts occur from the analysis for a given transcript. Furthermore, we ensure that the nucleotide sequence expected as the reference allele for each variant is present in the canonical cDNA before substituting it for the alternative allele. Haplotypes where a variant presents such inconsistency are removed from further analysis. However, this conflict has not occurred using the data set by 1000 Genomes Project [10].

### Reducing the database size

As our goal is to remove unlikely protein sequences, we provide a number of optional pre- and post-processing steps. Transcripts can be filtered to include only those of a desired biotype (e.g., protein-coding transcripts) as annotated in Ensembl. Alternatively, the user can provide a list of transcript identifiers that are to be included. Moreover, transcripts lacking the annotation of the start codon position can be discarded. Variants are filtered to a user-defined minor allele frequency (MAF) threshold. After the database is created as described above, sequences that match only to translations of untranslated regions (UTRs) are removed by default (Figure 4C). The user has the option to keep the translations of UTRs.

In protein haplotypes differing from the reference sequence, on average, five variants in linkage disequilibrium (LD) present the alternative allele. However, even if we only consider allele combinations found in the data set of phased genotypes, we encounter a substantial number of infrequent combinations. Removing haplotypes with a frequency below a certain threshold from the database can inadvertently render common alleles invisible in cases where these common alleles are dispersed across a multitude of rare haplotypes. Therefore, we have, by default, disabled the thresholding of haplotypes by frequency, preserving the breadth of genetic diversity within the database. However, we provide users with the flexibility to apply their own thresholding criteria, whether by frequency or by the number of occurrences, to suit the specific demands of their research.

It is worth noting that this issue becomes particularly pronounced when considering variants that map to the untranslated regions (UTR) of spliced transcripts. As these variants are both common and numerous, they contribute to a high number of haplotypes. Consequently, the inclusion of variants mapping to UTRs is, by default, disabled, offering users the flexibility to enable this feature when desired. This adaptability ensures that researchers can tailor the database to their specific analytical needs, striking a balance between inclusivity and precision in their investigations.

### Classification of peptides

We classify the peptides that can be matched to sequences generated by ProHap into four categories (Figure 5) as defined in our previous study [11]: canonical peptides map to *canonical* protein sequences. *Single-variant* peptides contain the product of the alternative allele for one genetic variant. *Multi-variant* peptides contain the product of the alternative allele for two or more genetic variants. Moreover, we introduce a fourth category - *frameshift* peptides map to protein sequences containing a frameshift mutation, or occurring downstream of frameshift and upstream of the early stop codon.

**Figure 5:**
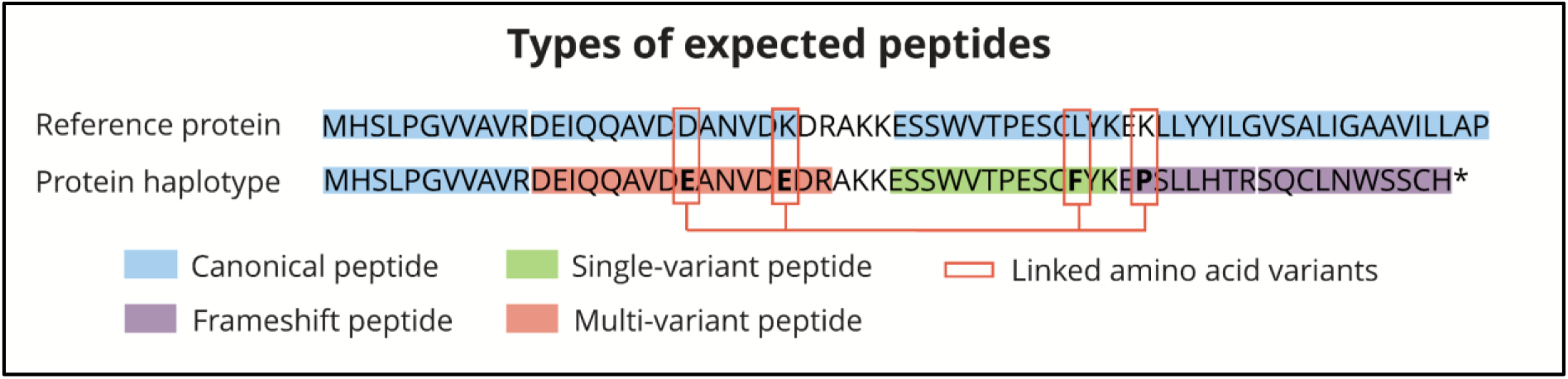
Example of a protein haplotype including four peptide categories that are expected. Peptides are marked by the colour representing the category, amino acid substitutions resulting from the alternative allele of three co-occurring genetic variants are marked by red squares. The asterisk sign (*) represents a stop codon.

If a peptide is canonical with respect to one protein sequence, and single- or multi-variant with respect to another protein sequence, it is considered a canonical peptide. Similarly, if a peptide is single-variant with respect to one protein sequence and multi-variant with respect to another protein sequence, it is considered a single-variant peptide. Peptides matching to contaminant sequences are discarded from further analysis.

We provide a Python script along with ProHap, which will classify the peptides provided in a list of peptide-spectrum matches, e.g., as an output of PeptideShaker [18], given the output table and FASTA file by ProHap.

We consider a variant to be discoverable if it satisfies the following conditions: (a) the protein consequence can be identified in a single-variant, multi-variant, or frameshift peptide of length between six and 40 amino acids with up to two missed cleavage sites (four for chymotrypsin), and (b) the peptide does not map to a canonical or contaminant sequence.

## Data Availability

The six databases of protein sequences derived from the phased genotypes of the 1000 Genomes Project, along with all necessary metadata, are available at https://doi.org/10.5281/zenodo.10149278

## Code Availability

ProHap’s source code, along with all documentation, is available at https://github.com/ProGenNo/ProHap. The pipeline used downstream of ProHap to obtain the results presented in this study is available at https://github.com/ProGenNo/ProHapDatabaseAnalysis.

## Acknowledgements

This work was supported by the Research Council of Norway (project #301178 to MV), the University of Bergen. PRN was supported by grants from the European Research Council (AdG #293574), Stiftelsen Trond Mohn Foundation (Mohn Center of Diabetes Precision Medicine), the University of Bergen, Haukeland University Hospital, the Research Council of Norway (FRIPRO grant #240413), and the Novo Nordisk Foundation (grant #54741). LK was supported by the Wallenberg AI, Autonomous Systems and Software Program (WASP) funded by the Knut and Alice Wallenberg Foundation.

The computations were performed on the Norwegian Research and Education Cloud (NREC), using resources provided by the University of Bergen and the University of Oslo. https://www.nrec.no

This research was funded, in whole or in part, by the Research Council of Norway 301178. A CC BY or equivalent license is applied to any Author Accepted Manuscript (AAM) version arising from this submission, in accordance with the grant’s open access conditions.

## Competing interests

The authors declare no competing interests.

## Notes

https://doi.org/10.5281/zenodo.10149278

### Competing Interest Statement

The authors have declared no competing interest.

### Summary of Updates

The changes in the manuscript are the following: 1. We explained the term "superpopulation" as used in the 1000 Genomes data set. 2. We added a few explanatory labels to Figure 1 and got rid of some of the abbreviations. 3. We have created six different protein databases, thresholding variants at 1% minor allele frequency within each of the five superpopulations separately, as well as globally for the complete database. This change in thresholding influenced the statistics reported in Figure 2 and the preceding paragraph, although all our conclusions still hold.

